# Spectral measure of color variation of black - orange - black (BOB) pattern in small parasitic wasps (Hymenoptera: Scelionidae), a statistical approach

**DOI:** 10.1101/652594

**Authors:** Rebeca Mora, Marcela Hernandez-Jimenez, Marcela Alfaro-Córdoba, Esteban Avendaño-Soto, Paul Hanson

## Abstract

A group of eight scelionid genera were studied by means of microspectrophotometric measurements for the first time. The orange and black colors were analyzed quantitatively, which in combination with Functional Data Analysis and statistical analysis of Euclidean distances for color components, describe and test the color differences between genera. The data analyzed by means of Functional Data Analysis proved to be a better method to treat the reflectance data because it gave a better representation of the physical information. When comparing the differences between curves of the same color but different genera, maximum differences were present in different ranges of the spectra, depending on the genus. Reflectance spectra were separated into their spectral color components contributions (red, blue and green). Each component had its own dominant wavelength at the maximum of the spectrum. We found differences in the dominant wavelength for specimens of the same genus, which are equivalent to differences in the hue. A correlation between the mean values of characteristics of the color components was used in an attempt to group the genera that show similar values. The spectral blue components of the orange and black areas were almost identical, suggesting that there is a common compound for the pigments. The results also suggest that cuticle from different genera, but with the same color (black vs black, orange vs orange) might have a similar chemical composition.

## Introduction

Light conveys information and plays a fundamental role in biology, providing the ability to extract useful information from the environment which is crucial for the survival of many insect species. Furthermore, light and other specific features from the visual world allow organisms to recognize one another and cope with their surroundings.

The reflection, absorption or scattering of specific wavelengths of light by surface structures, the deposition of different chemical pigments in the outer body layers or the interaction between various mechanisms, such as pigments amplifying iridescence in butterflies [1], produce a remarkable variety of colors with complex and diverse patterns in insects. These markings contribute to important behavioral, neurological, physiological, visual, morphological and developmental traits [2–9].

Compared with Lepidoptera and Coleoptera, there are relatively few studies of color patterns in the order Hymenoptera. What studies exist are mostly related to the yellow and black pattern in Vespidae [10, 11] and the variation in color patterns in *Bombus* spp. [12]. In Hymenoptera there are also few studies regarding the chemical and physical nature of the color patterns exhibited, some of the pigments reported being pheomelanins (black, orange-red, yellow variations in velvet ants and bumble bees) [2, 13], melanin (brown-colored cuticle) and xanthopterin (yellow-colored cuticle) in *Vespa orientalis* [14].

Scelionids show a recurring color pattern of black head, orange mesosoma, and black metasoma (BOB pattern; Fig 1) which has also been documented in species of numerous other hymenopteran families [15]. The wide distribution of the BOB pattern in Hymenoptera suggests that the coloration arose due to changes in shared metabolic or genetic routes as reported in other taxa, and strongly suggests that it has some biological function [10].

**Fig 1.**
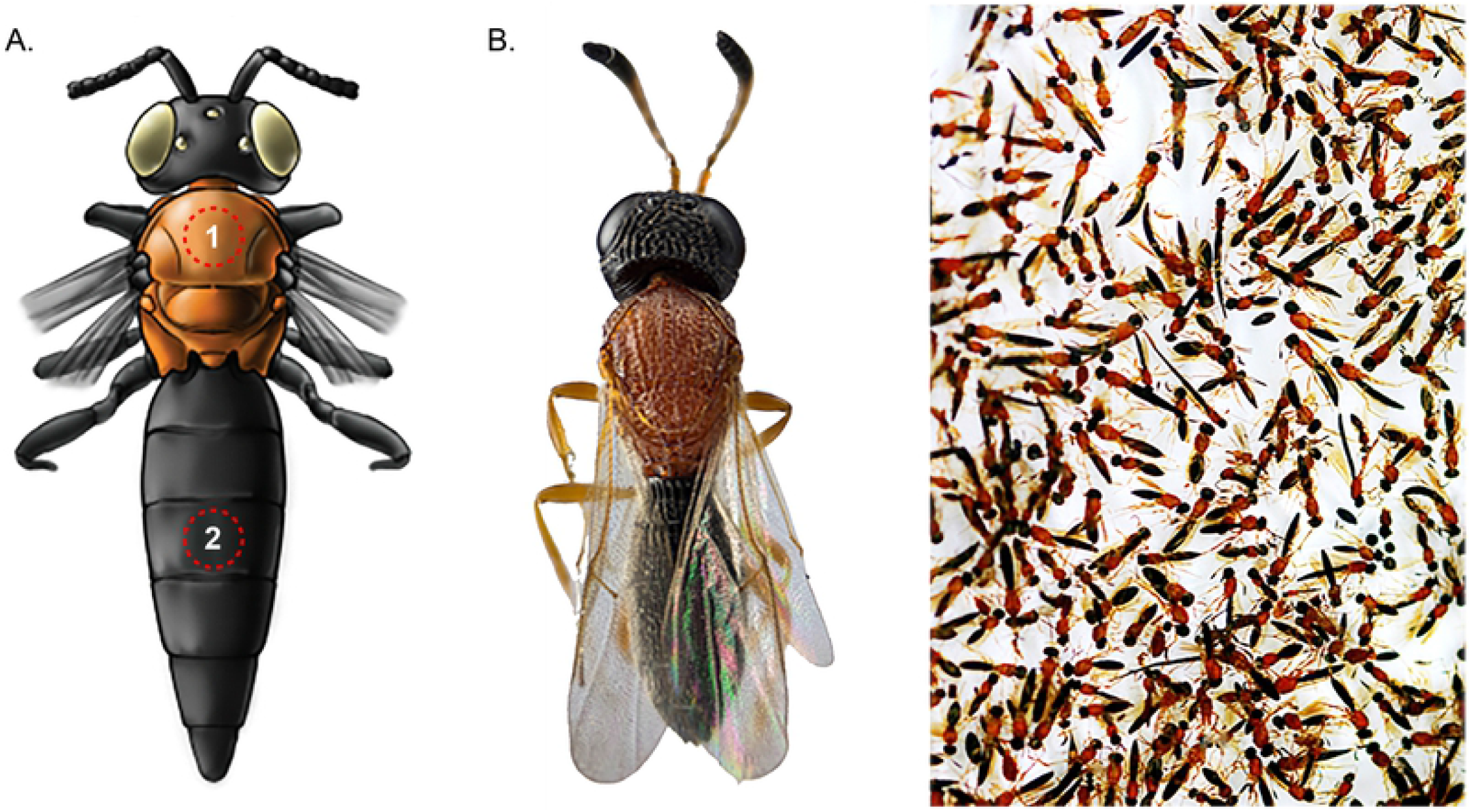
BOB Pattern Illustration. (A) Detail of the area (highlighted square) used for measurements of both colors, zone of 1 mm2 of mesoscotum (1) and third tergite of metasoma (2). (B) This figure also shows frozen insects recently brought from the field with the BOB color pattern. The focus stacked macro photography (specimen of 0, 6 mm) was obtained with a Reflex 850 camera coupled to a 20x microscope lens. The photograph of field insects was taken with a Canon SX10 IS camera with a 105mm macro lens.

It has been reported that most animals perceive color differently than humans [16] which has created a new surge of interest motivating widespread efforts to quantify both color traits and their visual environments. This new focus on generating objective information about the coloration of insects provides a more functional and evolutionary perspective, giving us a new tool to understand the real magnitude of diversity of color in insects.

Color characterization of specimens, especially in taxonomy, has been done mostly subjectively, although more quantitative methods have been proposed [17]. There are few reports treating the measurement of color rigorously or analyzing spectral data and its variability as a functional measurement, which potentially can show differences detected by insects, and shed a light on their biology.

Among many studies that measure and compare colors in insects, the methods vary widely, see for example [18], [19], [11]. Endler et al. [20] take into account the continuous nature of the wavelengths as measures of color and use Principal Component Analysis (PCA) to analyze patterns of color, while recognizing its limitations. Functional data analysis (FDA) is a powerful method to study spectrally measured data [21]. For example, Rivas et al. [22] propose the use of FDA techniques to test color changes in stone, and present a comparison with a multivariate method.

A statistical approach of reflectance measurements of the dorsal surface of eight Scelionidae genera was used in order to study the BOB color pattern. The reflectance spectra were measured by means of a microspectrophotometer. A statistical description of the BOB color pattern was obtained by means of different statistical approaches, applied to the spectral information as a function of genus, color, spots of measurement and specimens.This approach was used due to the small size of these wasps and the difficulty involved in obtaining sufficient fresh material for chemical analysis of the pigments.

## Materials and Methods

### Microspectrophotometry (MSP)

The reflectance spectra of 8 genera belonging to the Hymenopteran family Scelionidae were measured with a 508 PV UV-Visible-NIR Microscope Spectrophotometer (MSP) (CRAIC, Los Angeles, USA). The MSP is coupled to an Eclipse LV100 ND Microscope using episcopic illumination (Nikon, Tokyo, Japan). A spot size area of 200*μ*m x 200*μ*m was measured. A spectralon standard was used as a white reference because of the diffuse reflection produced by the roughness of the surface of the cuticle. Table 1 includes further information regarding the measurement settings.

**Table 1.**
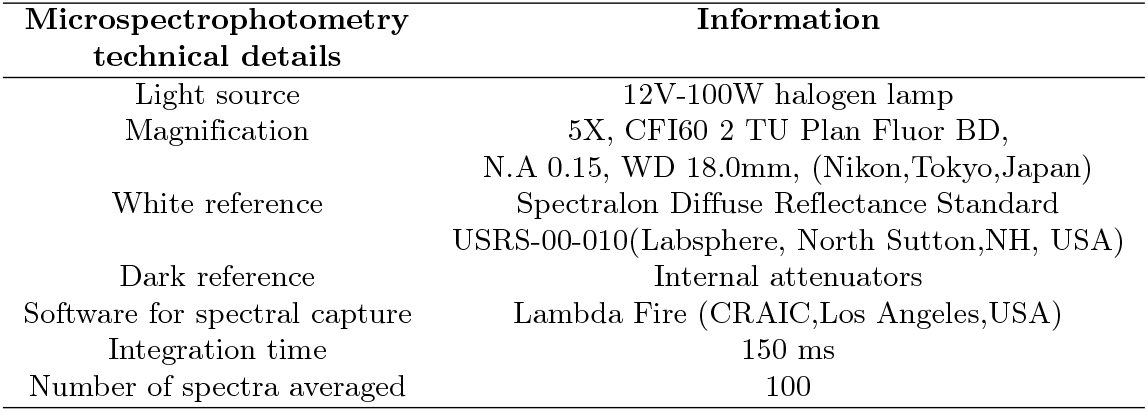
Information about the analysis of color data.

### Specimens and sampling for optical measurements

Recently collected specimens are crucial to ensure the good condition of the external cuticle structures. Since Scelionidae are small in size and their hosts (insect eggs) are difficult to find, collecting specimens requires considerable effort.

Specimens of 8 genera were collected: *Acanthoscelio*(AC), *Baryconus*(BA), *Chromoteleia*(CR), *Lapitha*(LA), *Macroteleia*(MA), *Scelio*(SC), *Sceliomorpha*(SM), *Triteleia* (TR). They were found and caught with entomological nets over a period of 12 months. Additionally some specimens of the genus *Evaniidae*(EV) were included, because they exhibit a similar BOB pattern, but are quite far removed from scelionids in the phylogeny of Hymenoptera [23].

The collection site was a patch within the protected zone of El Rodeo, which is part of the Mora Municipality, located 30 Km southwest of San José, Costa Rica. This natural reserve is composed of a secondary forest and the last remnant of primary forest (approximately 200 ha). Specimens were identified by R.M. and P.H. and voucher specimens were deposited in the Museum of Zoology of the University of Costa Rica. Due to the lack of taxonomic studies, species level identifications were not possible for most of the included genera, but by collecting from just one locality the problem of mixing species within a genus should be minimal (though not completely eliminated). A previous study demonstrated that males and females do not differ in color [15].

All experimental protocols were carried out in accordance with the involved institution’s guidelines and regulations. The specimens were killed by freezing and then, by using a needle cut and polished on both ends, one square millimeter or a small notch of area was dissected in the scutum of the mesosoma (orange) and in the third tergite of the metasoma (black) (Fig 2). The orange and black samples of 4 specimens of each of the 8 genera were analyzed. For each specimen three samples were taken, from the mesosoma (orange) and from the metasoma (black).

**Fig 2.**
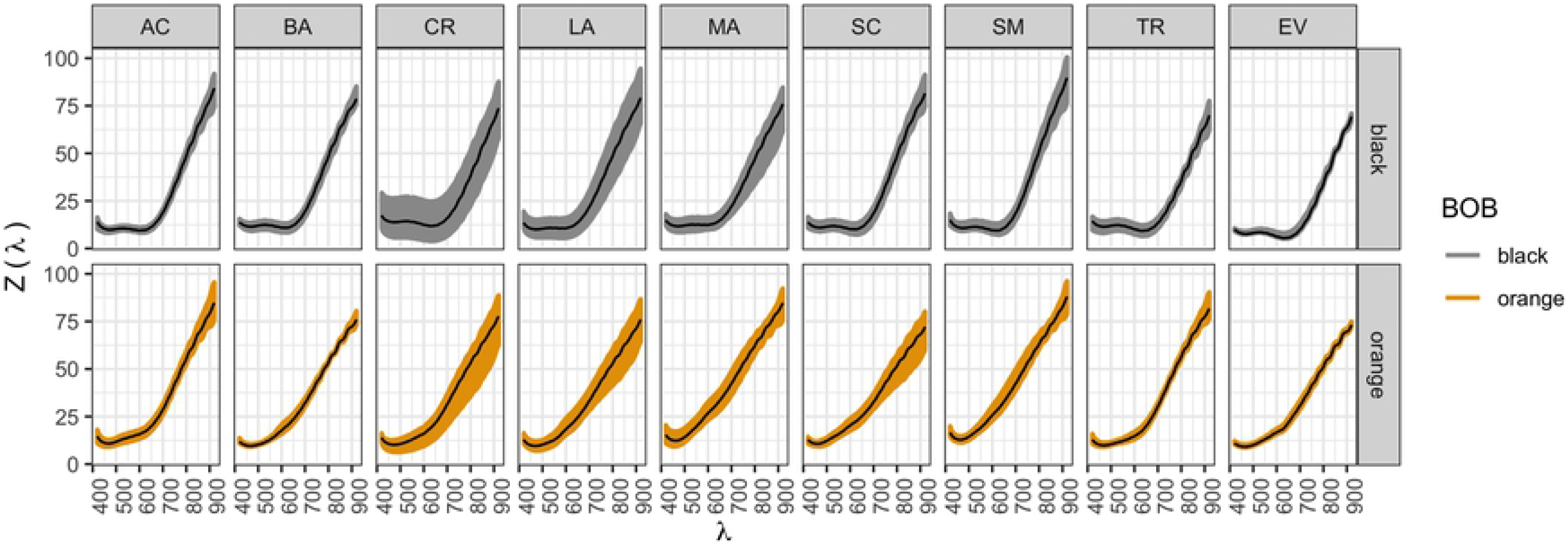
Reflection functions. Functions *Z*(*λ*) in y axis, for each color within each genus. Black lines show the functional mean per case.

### Colorimetric differences

Fig 2 shows the spectral reflectance of the eight genera for the black and orange segments together with the control genus. The data are presented with the average reflectance in black and the rough data showing the dispersion in gray and orange.

By using the measured spectra (Fig 2), the CIE-XYZ tristimulus values were calculated (see S1). A D65 day light standard illuminant for the CIE observer was chosen. Following the CIE recommendations, a spacing of 1 nm for wavelength was used. Since the data measured with the microspectrophotometer have a different wavelength spacing, an interpolation was made using the splinefun function in R [24], in order to generate a list of spectral points with the desired spacing. After the tristimulus values were obtained, they were used to calculate the CIE 1976 (L* a* b*) (CIELAB) color space coordinates, using the corresponding definitions as detailed in Appendix S1.

The CIELAB color space is Euclidean, therefore distances between points can be used to represent approximately the perceived magnitude of color differences between object color stimuli of the same size and shape, viewed under similar conditions. The CIE 1976 L, a, b (CIELAB) color difference definition was adopted [25]:

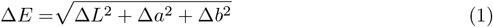

Here, the quantities Δ*P*, with *P* = *L, a* or *b*, represent the subtraction of the corresponding coordinates of the two colors being compared. There are other CIE definitions for color differences, see for example the ones explained in reference [25]. Nevertheless, as discussed by Melgosa [26], the definition in Eq 1 yields results that are not very different from more recent and complex definitions. Since the purpose of this work is not to evaluate the color space metrics, but to consolidate a quantitative framework to compare colors using the information available from MSP, the CIE 1976 definition (Eq 1) was chosen for the sake of simplicity. Moreover, regarding the definition of a color difference threshold, it can be argued that CIE color spaces are based on human vision sensibility, and the color difference threshold will depend not only on the spectral region but also on the observer. Nevertheless, in order to establish a mean threshold value to define whether two colors are different, we used the value 2.3 as reported by [27] specifically for CIELAB. In this sense, when the color difference Δ*E* is less than this value, colors are considered as equal. Otherwise, colors are said to be different.

#### RGB spectral components

As a way to access information about similarities and differences between the reflectance spectra in different portions of the visible electromagnetic spectrum, a thricromatic decomposition is proposed using the CIE color matching functions (CMFs) as follows:

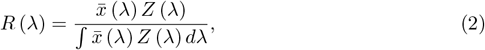

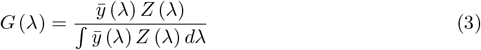

and

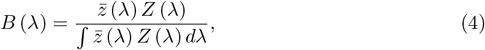

where *Z*(*λ*) is the measured reflectance, 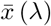, *ȳ*(*λ*) and 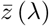 are the CIE 1976 CMF’s. [25].

### Statistical Analysis

#### Data sets

Each of the observations has 1185 recorded points: the reflectance (*Y*) as a function of wavelength value (*λ*), where *λ* is a sequence of 1185 equidistant points from 420.2 nm to 919.7 nm. The use of (*λ*) instead of the usual (*s*) in FDA, is done to match the representation of a wavelength in the physics literature. For the remainder of this paper, the 1185 points will be referred as reflectance curves, which were recorded in the following way: each curve was constructed with the reflectance of color in mesoscutum spots per colored area, and was measured three times in each specimen, in different spots. The observations correspond to the reflectance measured in the orange mesoscutum areas versus the black metasoma areas for 36 specimens: 4 specimens for each of the 8 genera, and 4 specimens for *Evaniella*, which was observed as a control. The response variable is either a difference of reflectance or a measure of reflectance, and it can be measured in three different levels:

- The univariate mean measure: *m_ijk_* where *m* is the mean difference between black and orange measures for all possible wavelengths in curve *ijk*, where *k* represents repetitions, *j* specimens, and *i* genera. This is is used as a naive base comparison.
- The functional measure: *Y_ijk_*(*λ*) where the difference in reflectance between black and orange curves *Z*(*λ*) is represented by *Y*, and was measured for each wavelength *λ*, and repeated *k* times in each specimen *j*, and genera *i*. In this case, *Y_ijk_*(*λ*) describes a reflectance difference curve.
- The multivariate distance measure: a Euclidean distance Δ*E* as defined in 1 for each pair of curves being compared.

A table was created relating the different factors (spots, specimen, and genera) with the response in the univariate and functional case. In total, 96 combinations of 3 spots, 4 specimens, and 8 genera were recorded. In the multivariate distance case, a total of 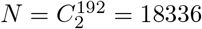 comparisons were used as observations, and then it was recorded whether or not each pair had a difference: genus, color, spot and specimen.

#### Data Analysis

Prior to the analysis, descriptive statistics were generated for each genus and color section (black and orange). The objective of the data analysis is to contrast the null hypothesis of no difference between means of each combination of genus and color area, and for that, univariate and functional ANOVA were performed. The multivariate distance case was also analyzed to match the physics literature on color differences. The following methods were used in each case:

- Univariate method: A one-way ANOVA was used to compare the mean difference between black and orange for all possible bandwidths *m_ijk_* with genus as the factor and controlling for specimen and spot.
- Functional method: A FANOVA was used to compare the difference curves *Y_ijk_*(*λ*) with genus as the factor, and controlling for specimen and spot. A statistical test was performed to interpret the contrast results for the mean difference curves. R packages erpFtest [28], and fdANOVA [29] were used.
- Multivariate distance method: A two-way ANOVA was used to compare the Euclidean distance Δ*E* between each combination of color and genus, using dichotomous variables as factors: color as 0 if the pair is from the same color, 1 if it differs; genus as 0 if the pair is from the same genus and 1 if it differs. Here also, specimen and spot were controlled.

The objective is to illustrate the functional method compared to the usual univariate practice, and with a multivariate distance alternative, to highlight the importance of taking into account all the information from the difference curves. More specifically, for the univariate and functional responses, the ANOVA model can be written as the following:

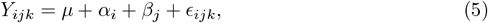

for *i* = 1, 2,..*p* = 8 species, *j* = 1, 2,.., *r* = 4 specimens, and *k* = 1, 2, 3 sampled spots in each colored area from each of the 4 specimens. Usual restrictions and assumptions for the ANOVA model apply:

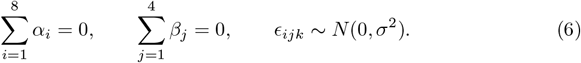

In the model, the global mean *μ* = 0 given that we are working with difference curves, and *α_i_* is the genera effect of level *i* and *β_j_* is a block effect to explain the intra-genus variance. In order to account for the dependence between spots from the same specimens (in the two colored areas), a random effect *η_k_* was tested, and different covariance structures for Σ_*η*_ were fitted (assuming dependence *η_k_* ~ *N*(0, Σ_*η*_) or independence between spots). Lastly, *ϵ_ijk_* is the residual accounting for the unexplained variation specific to the *k*th observation, *i*th genus and *j*th specimen. Depending on the response *Y_ijk_*, the residuals *ϵ_ijk_* are a number (univariate) or a curve (functional). Assumptions were tested for all the model options.

For the multivariate distances, the ANOVA model differed in the factors:

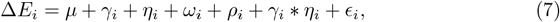

where the global mean *μ* represents the average change for the *N* comparisons, and then each of the factors add the change given that the pair has different genera (*γ_i_*), color (*η_i_*), spots (*ω_i_*) or specimens (*ρ_i_*) for each pair *i*. Finally, *ϵ_i_* is the error modeled as white noise. Assumptions were tested for this model as well.

Statistical analysis was performed using the computing environment R ([24]). The code and data to perform each of the tests mentioned in this paper are available for download in https://github.com/malfaro2/Mora_et_al.

## Results

The data provide support for reflectance difference variations between and within genera of the scelionids observed, when testing both the univariate data and the functional data (p-value < 0.00001 in all cases). The result is supported by the multivariate distance test, where there is evidence to say that overall change *μ* is greater than zero (p-value < 0.00001). Details about the significant factors that explain that change are summarized in the following sections.

### Overall differences

Fig 2 presents descriptive statistics for measured spectra for each genus and each of the spectral colors. There are some genera with more variability, and there is a tendency to have low reflectance at lower wavelengths for both black and orange spots. It is important to point out that the measurements were performed grouping genera and not species, a factor that may account for the variability in some cases such as *Chromoteleia*. Moreover, the effect of topography of the cuticle and homogeneity of the pigment distribution can partly explain the data dispersion in some of the genera.

The difference reflectance of the curves defined as: *Y_ijk_*(*λ*) are plotted in Fig 3. The patterns from Fig 2 appear more evident. For each genus, the difference curve between black and orange is around zero before wavelength 550 nm (spectra are similar), and has a different pattern of dissimilarities in the range between 550 nm and 750 nm, and the region between 750 nm and 900 nm. Thus the differences between colors in each genus can be explained depending on the wavelength range. This analysis cannot be done if only the univariate measures are studied, given the information reduction as presented in the supplementary material S2.

**Fig 3.**
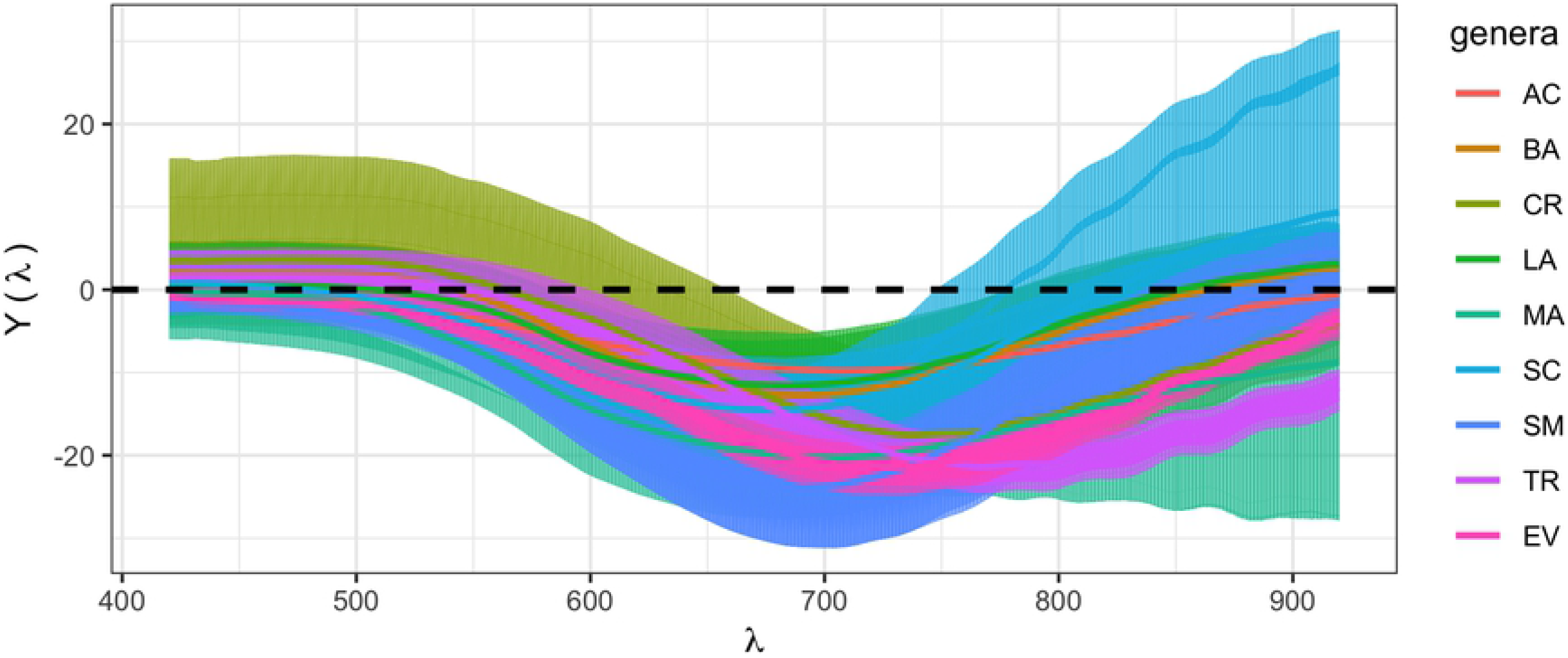
Difference of mean functions in each genus. Solid lines represent functional means, colored areas represent observations in each genus. A black dashed line represents the null hypothesis of no difference between colors in each genus.

In order to test the null hypothesis of *H*_0_: *α_i_* = 0 for all genera, models were used to fit the data for each response. The best fit in all cases was from the models that assume independence between spots, so linear models were used subsequently. The univariate (p-value < 0.0001) and functional (p-value < 0.0001) results coincide in that there is statistical evidence of a difference between mean reflectance of color for at least one genus and zero, controlling by specimen and spot. The following sections describe the genera that are significantly different from the rest.

The statistical tests confirm what the descriptive plots showed. For each genus, the difference between black and orange are significantly different from zero for both the univariate and functional case. Results from the functional ANOVA and a permutation test based on a Fourier basis function representation, to test whether there is a mean difference between colors and genera, show that the calculated statistic is *T* = 18.66961 [28] with an associated p-value < 0.0001, which means that there is evidence of at least one genera difference being different from the rest as shown in Fig 3.

The results for univariate comparison between genera can be found in Supplementary material S2. From such an analysis it can be concluded that the univariate approach is not suitable for treating this type of data.

### Multivariate Distance Analysis

ANOVA results for the model of Eq 7 presented in Table 2, show how the overall average change in all comparisons is 5.0535 units with a standard error of 0.2642, which makes the average statistically greater than the 2.3 threshold value. The most important factor in this case is when there is a change of color, since the estimate of *η* is 26.2081, which means that keeping all the other factors constant, the average Δ*E* is 26 units greater than the overall mean when the curves differ in color. Also, if the curves differ in both color and genus, the overall mean Δ*E* is 5.0535 + 4.3540 + 26.2081 +−4.3814 = 31.2342. The differences due to specimens are statistically not significant and the differences due to spots are on average 0.85, which makes it statistically significant but negligible on average, compared to the threshold.

**Table 2.** ANOVA Results for model 7.

|  | Estimate | Std. Error | t value | p-value |
| --- | --- | --- | --- | --- |
| $\mu$ | 5.0535 | 0.2642 | 19.12 | 0.0000 |
| $\gamma$ | 4.3540 | 0.2420 | 17.99 | 0.0000 |
| $\eta$ | 26.2081 | 0.3149 | 83.21 | 0.0000 |
| $\omega$ | 0.8524 | 0.1268 | 6.72 | 0.0000 |
| $\rho$ | -0.0425 | 0.1161 | -0.37 | 0.7142 |
| $\gamma:\eta$ | -4.3814 | 0.3358 | -13.05 | 0.0000 |

Given that it has been established that there is evidence of a difference between colors and genera in all analyses, there is a need to clarify which genera and which colors are making the difference. For that, a comparison matrix for Δ*E* was constructed and presented in two ways: comparisons between genera per color difference and comparisons between colors per genus difference. Such results are presented in Fig 4 and Fig 5 respectively. As an example, Fig 4 depicts how the distance Δ*E* between *Triteleia* (TR) and *Acanthoscelio* (AC) curves presents two distributions: one when both have the same color (in vermilion) and another when they have different colors (in bluish green). Both distributions present relatively low variability, and the first distribution is closer to zero, although its box - 50% of its data - is not close to zero or the threshold. In contrast, the distance Δ*E* between *Acanthoscelio* (AC) and *Acanthoscelio* (AC) curves for the same color (in vermilion) clearly include the threshold and can be taken as not significantly different.

**Fig 4.**
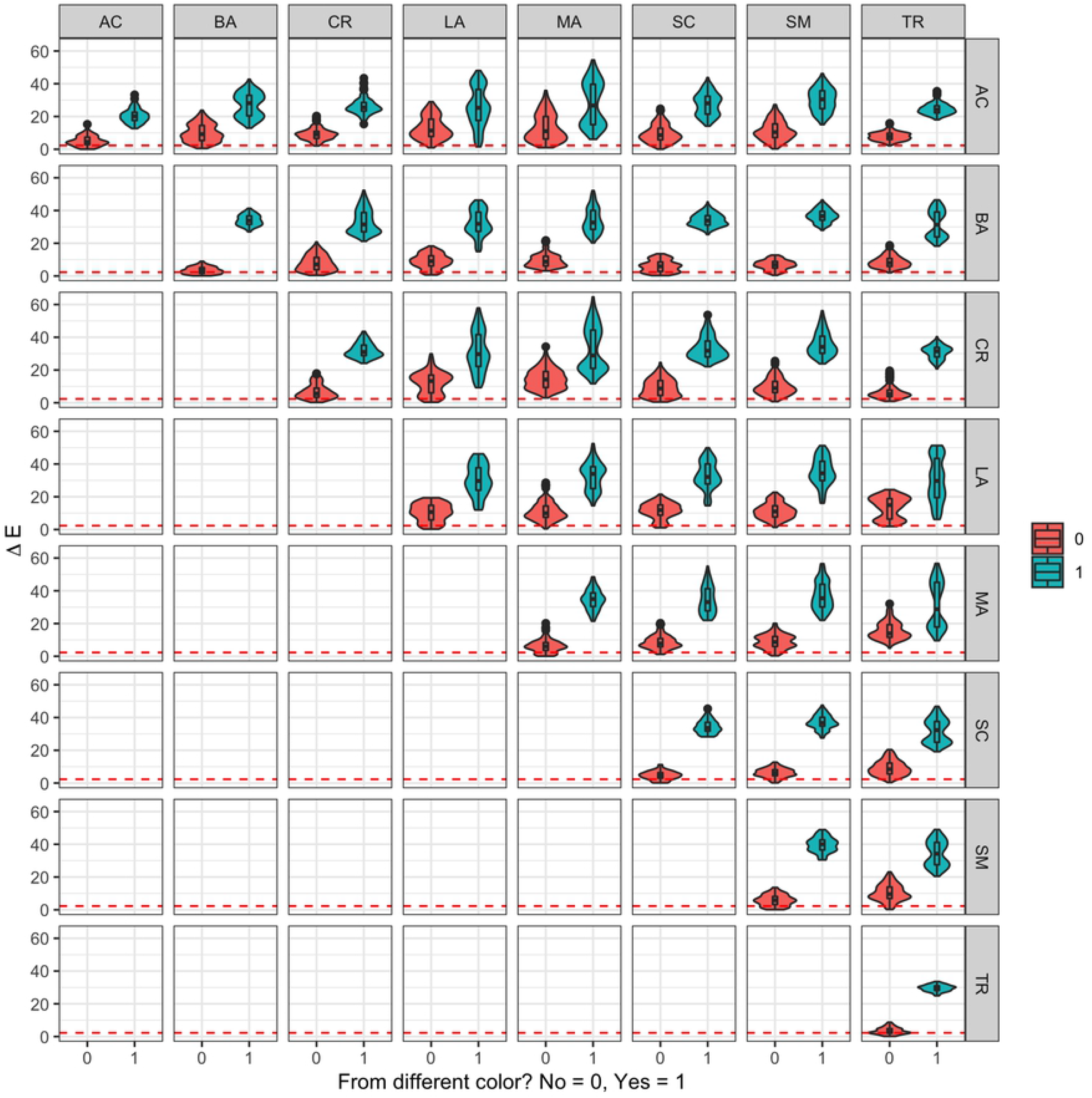
Violin and box plot comparison per genus. Distributions of Δ*E* calculated for pairs of curves of eight different genera: *Acanthoscelio* (AC), *Baryconus* (BA), *Chromoteleia* (CR), *Lapitha* (LA), *Macroteleia* (MA), *Scelio* (SC), and *Sceliomorpha* (SM), and two scenarios: when they have the same color (in vermilion) and when their colors differ (bluish green). The dashed vermilion line represents the threshold of 2.3.

**Fig 5.**
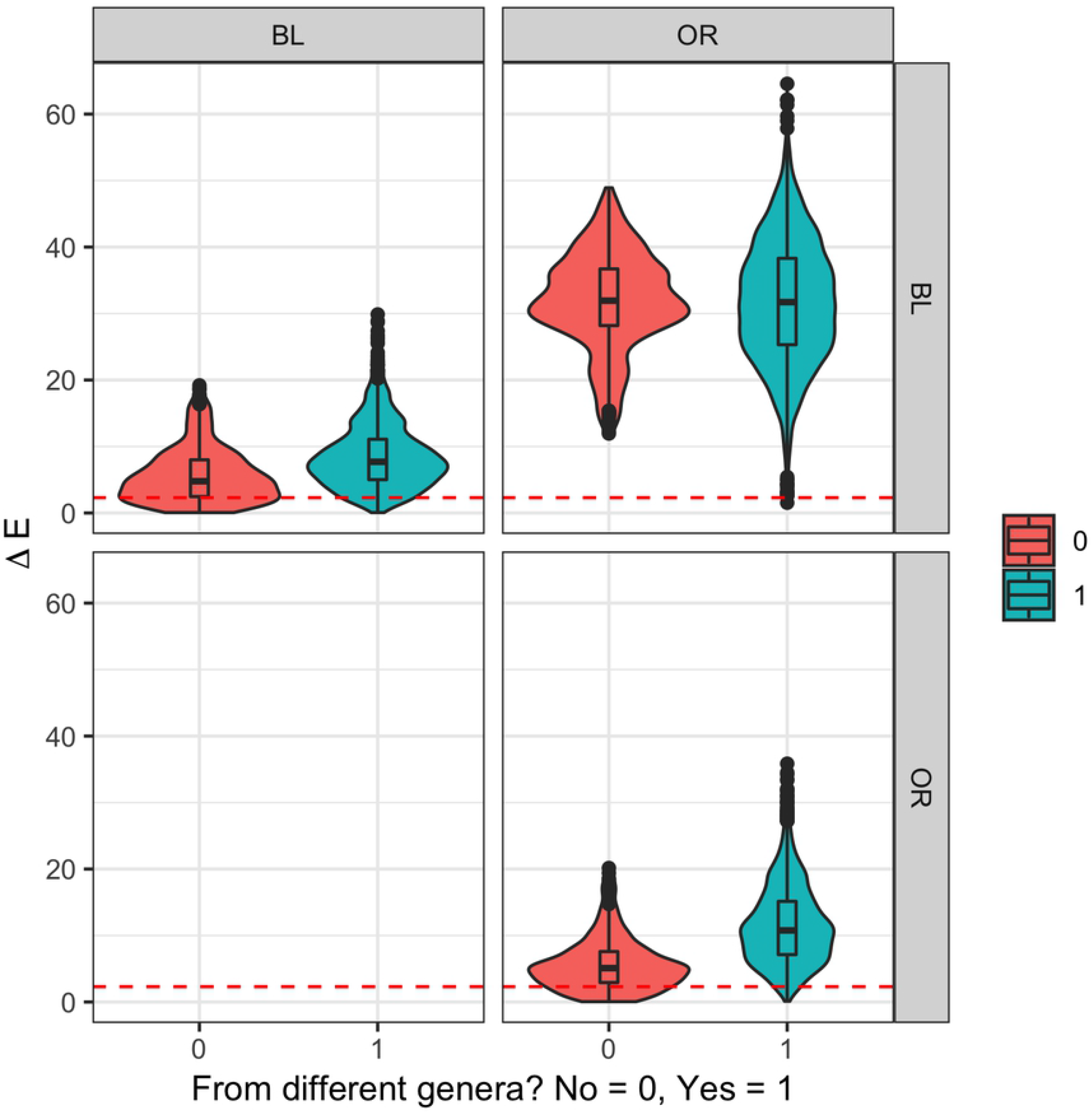
Violin and box plot comparison per color. Distributions of Δ*E* calculated for pairs of two different colors, black (BL) and orange (OR), and two scenarios: when they are the same genus (in vermilion) and when the genus differs (bluish green). The dashed vermilion line represents the threshold of 2.3.

The difference between distributions is not that clear when comparing colors. Fig 5 presents how in the case in which the colors differ (upper right corner) both distributions for Δ*E* are both clearly different from zero, while the distributions for Δ*E* of the curves with the same color are more difficult to distinguish from zero, especially if they are from the same genus (in vermilion).

The differences between curves can be described as an average of all Δ*E*, for each genus and color. By that metric, the maximum and minimum mean difference can be established, as described in Table 3 for comparisons between curves of different colors and Table 4 for comparisons between curves of the same color, pointing to the specific contrast with wider differences. Similarly, comparisons of different genera and the same genus are presented in S3 in the supplementary material.

**Table 3.** Top 10 according to 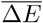 for comparisons of curves of the different color. Overall minimum mean is 2.96 and maximum mean is 45.08

| | Genus 1 | Genus 2 | Color 1 | Color 2 | $\overline{\Delta E}$ |
| --- | --- | --- | --- | --- | --- |
| 1 | CR | MA | BL | OR | 45.08 |
| 2 | MA | TR | OR | BL | 44.79 |
| 3 | MA | SM | OR | BL | 43.59 |
| 4 | LA | TR | OR | BL | 41.51 |
| 5 | CR | SM | BL | OR | 41.25 |
| 6 | SM | TR | OR | BL | 41.22 |
| 7 | MA | SC | OR | BL | 41.02 |
| 8 | CR | LA | BL | OR | 40.84 |
| 9 | SM | SM | BL | OR | 39.71 |
| 10 | BA | TR | OR | BL | 39.69 |

**Table 4.** Top 10 according to 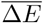 for comparisons of curves of the same color. Overall minimum mean is 2.96 and maximum mean is 45.08

| | Genus 1 | Genus 2 | Color 1 | Color 2 | $\overline{\Delta E}$ |
| --- | --- | --- | --- | --- | --- |
| 1 | AC | MA | OR | OR | 19.64 |
| 2 | MA | TR | OR | OR | 18.26 |
| 3 | CR | MA | OR | OR | 17.02 |
| 4 | AC | LA | OR | OR | 16.95 |
| 5 | AC | SM | OR | OR | 15.77 |
| 6 | AC | BA | OR | OR | 14.35 |
| 7 | LA | TR | BL | BL | 14.16 |
| 8 | SM | TR | OR | OR | 13.69 |
| 9 | LA | MA | OR | OR | 13.61 |
| 10 | CR | LA | BL | BL | 13.32 |

### RGB Spectral Components

Since it is difficult to point out why Δ*E* differs for each case, an extra analysis was performed taking the three color components of each curve: green, red and blue. The hypothesis in this case is that the differences that were found in terms of Δ*E* can be explained in terms of color components. For that, a similar exercise was done comparing color component curves: red (*R*(*λ*)), green (*G*(*λ*)) and blue (*B*(*λ*)), in terms of distance:

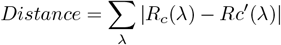

for each pair of comparisons (*c, c′*). The results in Fig 6 show how the same differences depicted when using Δ*E* are very clear for the green and red components, but not for the blue component.

**Fig 6.**
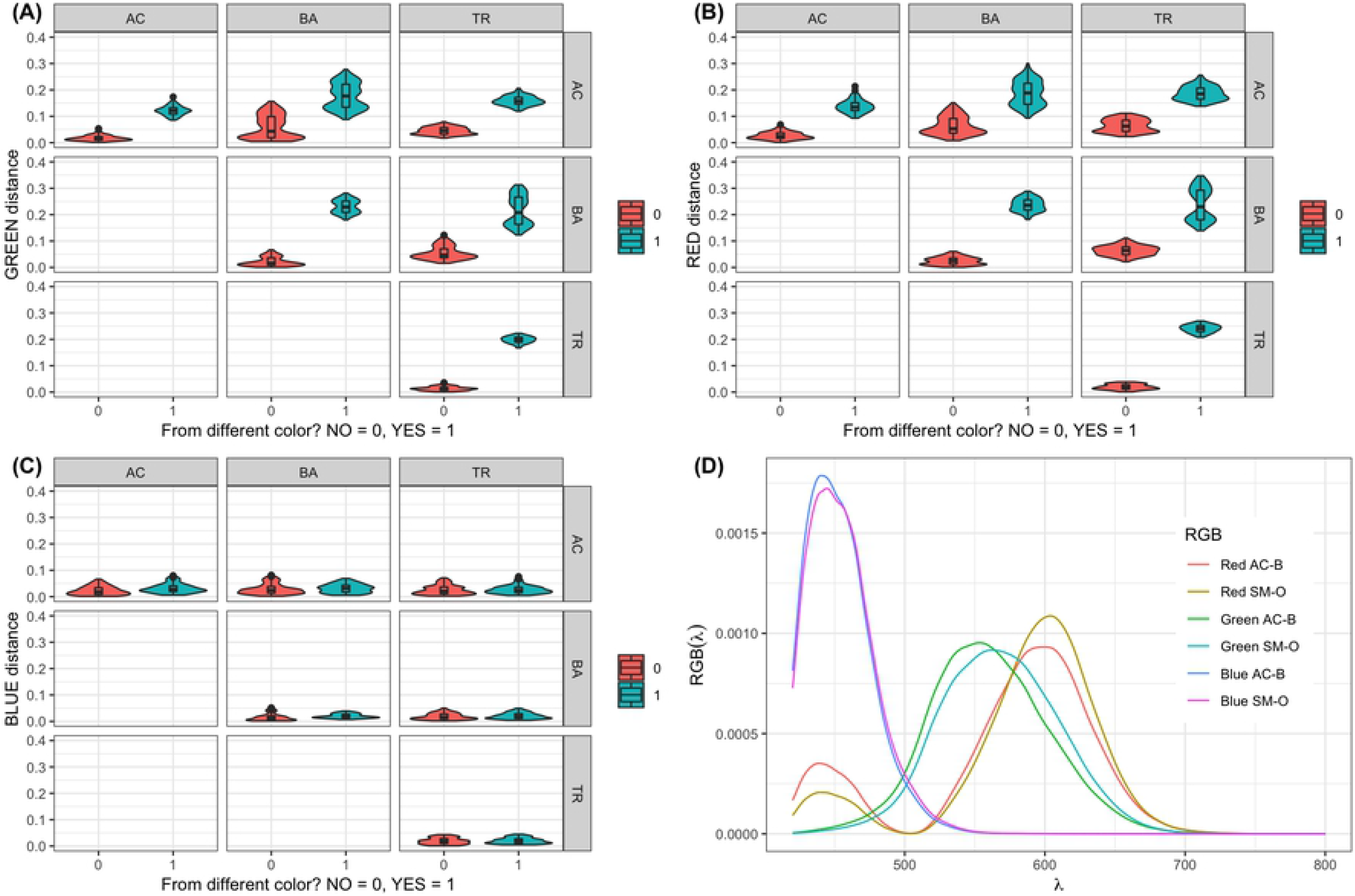
Examples of violin and box plot comparison per genus and color component. Distributions of Distance for (A) green, (B) red and (C) blue components, calculated for pairs of curves of three different genera: AC, BA, and TR, and two scenarios: when they have the same color (in vermilion) and when their colors differ (bluish green). Plot (D) is an example of how the R, G, and B curves look for AC in black spots and SM for orange spots.

Fig 6 in panel (D) shows the color component contributions for the AC and SM genera for the black and orange measurements. The spectra were normalised by its own area to illustrate the similarity of the contributions that are reflected in all data taken and presented in Fig 4 and 6 in panels A, B and C. Apart from the small changes in intensity for the normalised spectra, the maxima of each curve have a different dominant wavelength between genera, but the shape of the curves was very similar. The main changes are the differences in area when not normalised.

Our group has found evidence of a correlation between the color and/or reflectance spectra of beetles of the genus *Chrysina* (Scarabaeidae) with respect to its phylogenetic tree. Following the same line of thought, Fig reffig:F7 shows the scatter plots for the black and orange measurements for all data, representing the logarithm of the area of each color component versus the dominant wavelength at the maxima of each contribution. The mean is plotted as a solid circle of the same color for each set of faded data points.

The range of wavelength values at the maxima for each color component contribution is interesting, being around 5 nm for the blue contribution in both black and orange measurements. For the green (black measurement) and red (orange measurement) components it is about 5 nm, but for the red (black measurement) and green (orange measurement) it is approximately 20 nm.

The control genus *Evaniella* (EV) appears to separate from the group in the black measurements, but it is not that obvious for the orange measurements. Meanwhile, *Sceliomorpha* (SM) and *Macroteleia* (MA) group together for the green and red orange observations. The genus *Chromoteleia* (CR) is outside the main group in all three color components for the black observations and in the blue component of the orange observations. *Chromoteleia* (CR) and *Baryconus* (BA) also pair in the green and red components of the black and orange observations. Additionally *Sceliomorpha* (SM) and *Acanthoscelio* (AC) present a mean that is very close in the black measurements for the blue, green and red contributions. The genera *Lapitha* (LA) and *Triteleia* (TR) group together in the blue and green components for the orange contribution.

## Discussion

The differences between genera shown in Table 5 only take into account the aggregated data collected at all wavelengths considered. The difference in intra-genus variances illustrated in Fig 8 of the Supplementary material S2, confirms that the analysis of reflectance differences should not be done using univariate statistics. Said variability is due to the difference in measurements across wavelengths: some sections are equal to zero, others are statistically different from zero, as described in the previous section. This can yield misleading results, finding univariate groups statistically similar, when the functional results are describing something different.

**Table 5.** Mean reflectance *m_i_* for all possible bandwidths, by genus. Genera are ordered according to the difference magnitude. ** The univariate difference is not significantly different from zero. ^*a,b,c*^ Represent a group different from the rest, when doing univariate multiple comparisons. All analyses use *α* = 0.05. Color representation corresponds to the CIE 1976 (L* a* b*) color space coordinates calculated from the mean reflectance curves from Fig 1.

| Genera | Black | Orange | Absolute Difference |
| --- | --- | --- | --- |
| *Baryconus* | 29.5307 | 33.692 | 4.1613<sup>**,a</sup> |
| *Lapitha* | 29.5492 | 33.7317 | 4.1825<sup>**,a</sup> |
| *Scelio* | 29.3678 | 33.8015 | 4.4337<sup>**,a</sup> |
| *Acanthoscelio* | 29.2331 | 34.3834 | 5.1503<sup>a</sup> |
| *Chromoteleia* | 26.5133 | 33.3825 | 6.8692<sup>a,c</sup> |
| *Triteleia* | 24.0089 | 34.1785 | 10.1696<sup>b,c</sup> |
| *Sceliomorpha* | 30.683 | 41.7377 | 11.0547<sup>b</sup> |
| *Macroteleia* | 29.5722 | 41.3887 | 11.8165<sup>b</sup> |
| *Evaniella* | 21.2445 | 32.3144 | 11.0699<sup>b</sup> |

Functional Data Analysis (FDA) provides methods for analyzing data that are believed to arise from curves evaluated at a finite grid of points [21]. The present study highlights the differences in terms of results that could arise using univariate methods, which ignore physical information contained in the reflectance measurement as a function of wavelength, versus analyzing data using FDA techniques. In addition, a multivariate Euclidean distance defined in terms of a quantity with physical meaning such as Δ*E*, provides more evidence for analysis, besides confirming the FDA results. The details about multiple comparisons can be even more specific, if the color components are separated for each curve and the analysis is repeated.

The component analysis showed that the nature of the differences found using the FDA and Δ*E* methods are related to the green and red color components, but no correlation was found for the blue component. This is consistent with results obtained qualitatively using the difference curves *Y_ijk_*(*λ*) shown in Fig 3. Moreover, ANOVA results for model 7 (Table 2) confirm the reproducibility of the experimental protocol and, for most of the cases studied, the results suggest that there is a homogeneity of the sampling even though the taxonomic identification to just the genus level means that at least some of the genera could consist of more than one species.

This particular method for statistically testing for color differences differs from those in the literature due to the level of scrutiny. It goes from the general question, “Is there a difference between colors?” to more specific contrasts, using both the FDA and the color components methods. In this way, we are statistically comparing each pair of colors and genera, controlling for the other experimental factors.

In general, the results of the spectral decomposition, together with the color differences as analyzed in Fig 5, suggest that the cuticle segments from different genera, but with the same color (black vs. black, orange vs. orange) might have a similar chemical composition or an identical chemical composition with different concentrations. Tonal variations could be related to different amounts of an individual pigment or a combination of similar polymers distributed in the cuticle.

Recent studies report that the color phenotype is clearly associated with the concentration of certain pigment forms, but there could be other relevant factors besides concentration related to the expression of color [30]. In the case of pigments of different color (black vs. orange), the spectral blue components remain almost identical, suggesting that there is a common compound for the pigments. Preliminary chemical characterization by means of infrared spectroscopy showed subtle differences between spectra, suggesting that the pigments differ only by the nature of the functional group attached to the molecule. However, our preliminary results could not identify the nature of the molecules because the spectra contain contributions from the pigment together with other components of the cuticle.

A correlation between spectral color and phylogenetic relationship in beetles of the genus *Chrysina* (Scarabaeidae) has been observed in ongoing research by our group. However, the same does not appear to be the case in Scelionidae. For example, the orange color of *Macroteleia* and *Sceliomorpha* group together for the green and red components (Fig. 7B) despite these genera not being closely related in the most recent phylogenetic reconstruction [31]. Among the few phylogenetic correlations detected was for *Evaniella*, which belongs to a completely different family (Evaniidae) and might therefore be expected to differ markedly from all the other genera. This indeed appears to be the case for its black color, but not its orange color (Fig. 7). This is an interesting result in that it suggests that the orange color shows more convergent evolution than does the black color. Yet, within Scelionidae there are more differences in the orange color than in the black (Table 4). The above results merit further investigation.

**Fig 7.**
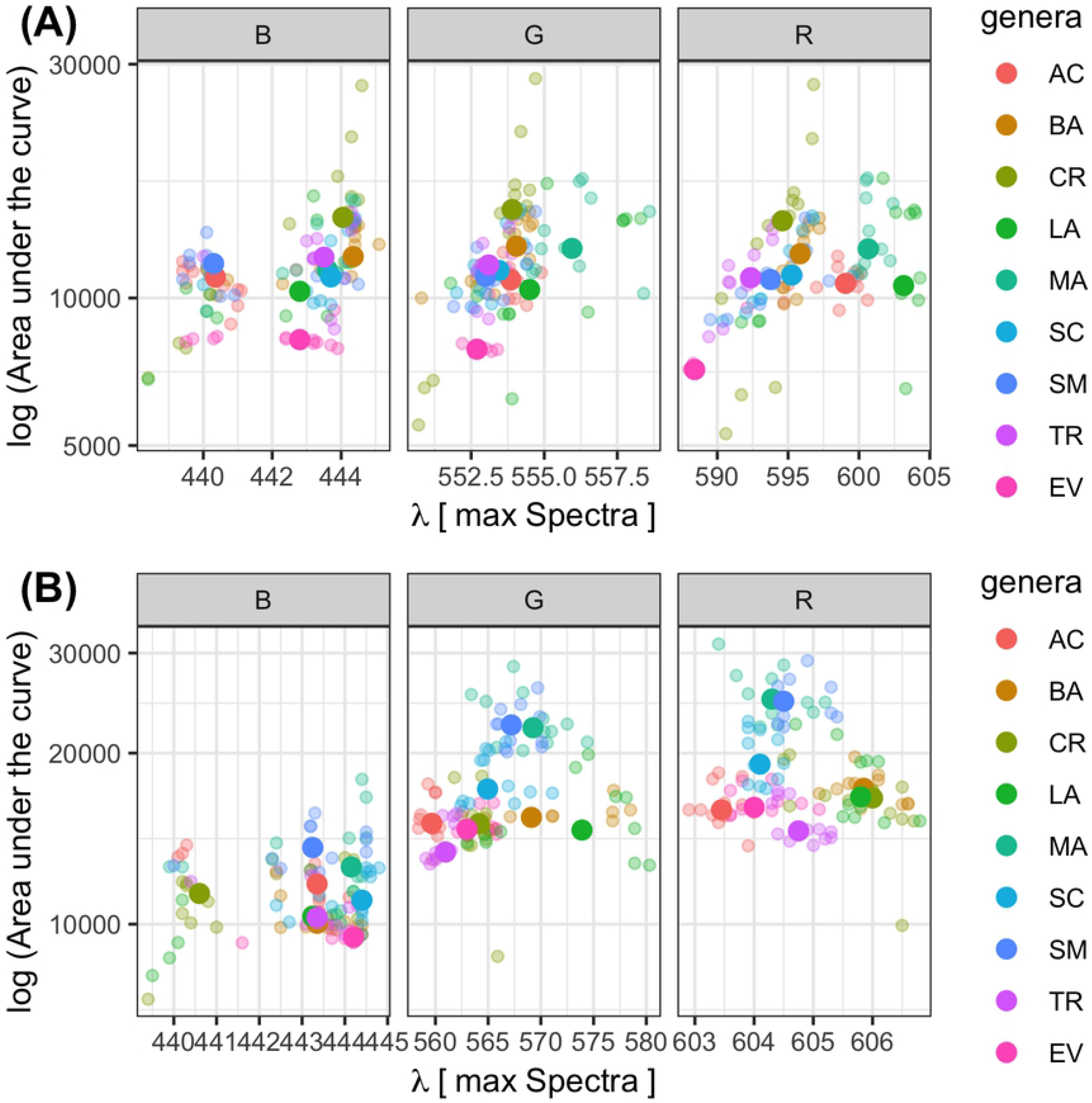
Scatter plot for color components per genus. Logarithm of the area under the curve, per component (red, green or blue) versus the corresponding wavelength in nm where the maximum occurs. Solid color points represent means for each genus and faded points correspond to independent observations. Plot (A) and (B) represent the components for the black and orange colors respectively.

In some other insects, contrasting black and orange color patterns are known to serve as aposematic (warning) coloration for potenital predators [32] and it is likely that the same is true of the BOB pattern, although this has not yet been tested. However, the orange mesosoma becomes less common at higher elevations [15], suggesting that other factors such as thermoregulation might play a role at these elevations. There is obviously much more to be learned about the black and orange colors present in numerous parasitic wasps, but it is hoped that the results of the present spectral analyses provide a framework for future research.

## Supporting information

### S1 Color coordinates calculation

The tristimulus values for color coordinates, according to the CIE (International Commission on Illumination, or Commission Internationale d’Eclairage) definition, were calculated as follows:

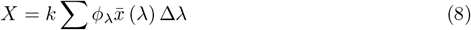

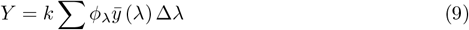

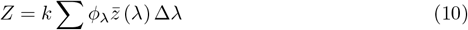

where 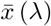, *ȳ*(*λ*) and 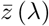 are the standardized color-matching functions (CMF) of the CIE 1931 standard observer. The constant k in equations 8, 9 and 10 is defined so that Y=100 for objects with perfect reflectivity. Finally, *ϕ_λ_* is the relative color stimulus function, defined as:

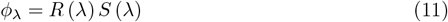

with *R*(*λ*)being the spectral reflectance factor (the measure spectra) and *S*(*λ*) is the relative spectral power distribution (SPD) of the illuminat. The coordinates in the CIELAB color space are:

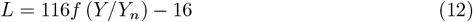

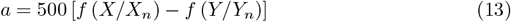

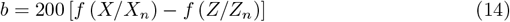

where

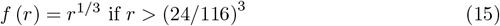

and

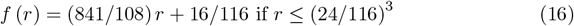

with *X_n_, Y_n_* and *Z_n_* being the tristimulus values of a idealized white reflector.

### S2 Univariate comparison for the inter-genera case

For the univariate case, Table 5 shows how *Baryconus, Lapitha*, and *Scelio* absolute differences are not significantly different from zero. This contradicts the functional data analysis result presented before, where all the difference curves are different between genera.

When describing the difference between genera, Table 5 shows how there are three groups of genera without univariate mean differences: group *a* consists of *Baryconus, Lapitha, Scelio, Acanthoscelio* and *Chromoteleia*, while group *b* consist of *Triteleia, Sceliomorpha, Macroteleia* and *Evaniella* as members. Lastly, *Acanthoscelio* and *Chromoteleia* (group *c*) seem not to differ in terms of univariate means. On the other hand, when examining the curves in Fig 3, it is clear that they differ, with *Triteleia*’s difference curve being very close to zero from *λ* = 420 nm to *λ* = 590 nm and then going to a negative difference of 20 around *λ* = 780 nm, while the curve for *Macroteleia* deviates from zero around *λ* = 500 nm and reaches −20 at *λ* = 680 nm. Given that other characteristics from the curve are compared in this case, it is reasonable to have a contradictory result. The pattern of contradictory results between univariate and functional results is repeated in all pairs of genera belonging to the same univariate group (a,b,c), which could indicate one of the limitations of the univariate analysis: it ignores the fact that the spectrometer measurements are curves and they should be compared as such.

Fig 8 shows the box plots for univariate mean differences for each genus. The black dashed line represents the null hypothesis of no difference between colors within a genus.

**Fig 8.**
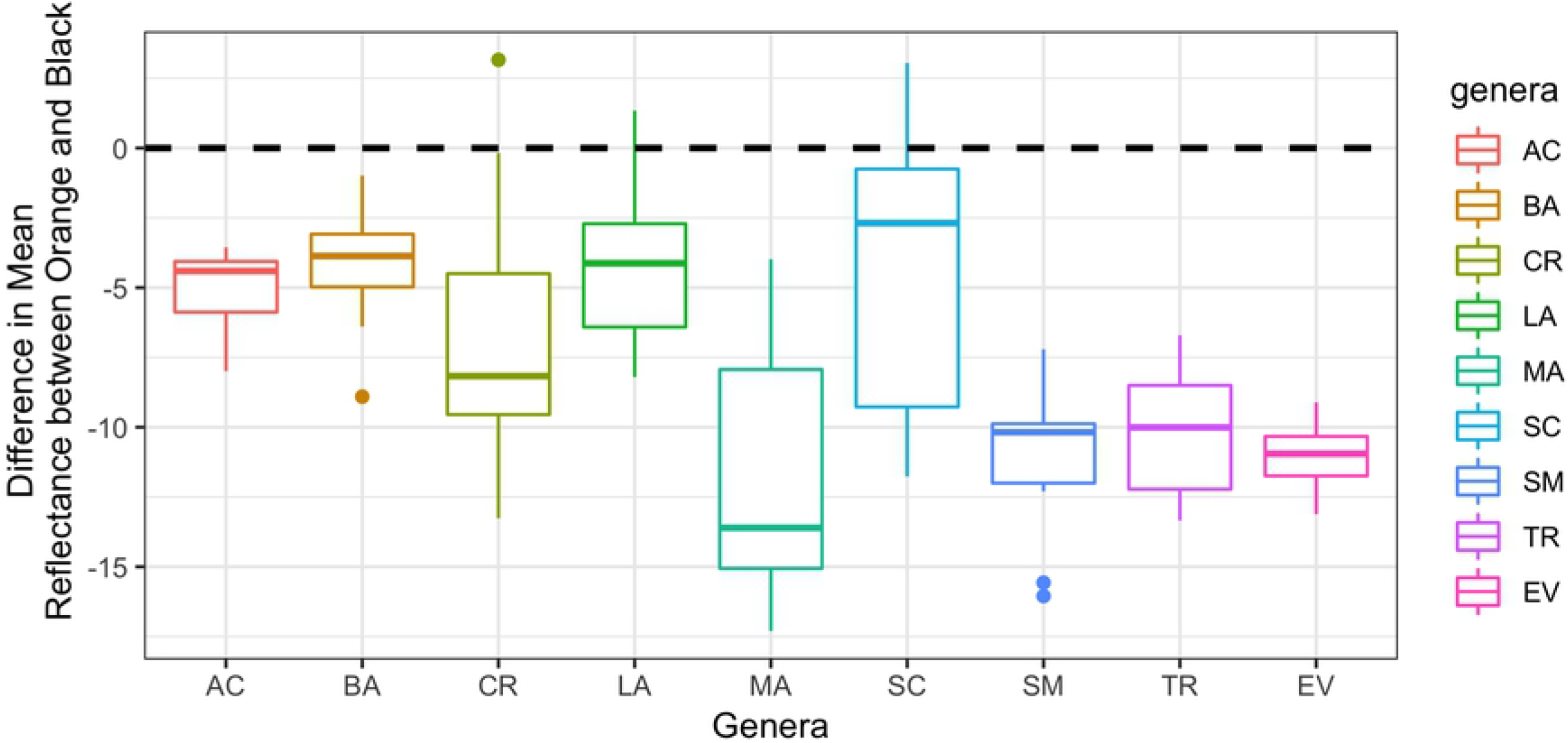
Box plots for univariate mean differences per genus. Black dashed line represents the null hypothesis of no difference between colors per genus.

### S3 Top 10 Table

**Table 6.** Top 10 according to 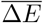 for comparisons of curves of different genera. Overall minimum mean is 2.96 and maximum mean is 45.08

| | Genera 1 | Genera 2 | Color 1 | Color 2 | $\overline{\Delta E}$ |
| --- | --- | --- | --- | --- | --- |
| 1 | CR | MA | BL | OR | 45.08 |
| 2 | MA | TR | OR | BL | 44.79 |
| 3 | MA | SM | OR | BL | 43.59 |
| 4 | LA | TR | OR | BL | 41.51 |
| 5 | CR | SM | BL | OR | 41.25 |
| 6 | SM | TR | OR | BL | 41.22 |
| 7 | MA | SC | OR | BL | 41.02 |
| 8 | CR | LA | BL | OR | 40.84 |
| 9 | BA | TR | OR | BL | 39.69 |
| 10 | AC | MA | BL | OR | 39.58 |

**Table 7.** Top 10 according to 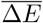 for comparisons of curves of the same genera. Overall minimum mean is 2.96 and maximum mean is 45.08

| | Genera 1 | Genera 2 | Color 1 | Color 2 | $\overline{\Delta E}$ |
| --- | --- | --- | --- | --- | --- |
| 1 | SM | SM | BL | OR | 39.71 |
| 2 | MA | MA | BL | OR | 34.59 |
| 3 | SC | SC | BL | OR | 34.36 |
| 4 | BA | BA | BL | OR | 34.11 |
| 5 | CR | CR | BL | OR | 32.02 |
| 6 | LA | LA | BL | OR | 30.26 |
| 7 | TR | TR | BL | OR | 29.57 |
| 8 | AC | AC | BL | OR | 20.37 |
| 9 | LA | LA | OR | OR | 10.40 |
| 10 | LA | LA | BL | BL | 10.28 |

## Acknowledgments

This work was inspired by personal conversations with Lubomir Masner. We would like to give special thanks to Mauricio Arce for the macro photography.

